# Systemic effects of cystic fibrosis transmembrane conductance regulator (CFTR) modulators on the blood proteome

**DOI:** 10.1101/2024.10.18.619058

**Authors:** Kerstin Fentker, Marieluise Kirchner, Matthias Ziehm, Sylvia Niquet, Oliver Popp, Julia Duerr, Laura Schaupp, Jobst Roehmel, Stephanie Thee, Susanne Hämmerling, Olaf Sommerburg, Mirjam Stahl, Simon Y. Graeber, Marcus A. Mall, Philipp Mertins

## Abstract

Cystic fibrosis (CF), resulting from a dysfunction in the cystic fibrosis transmembrane conductance regulator (CFTR), affects multiple organs through mucus obstruction and differences in secretion. The CFTR modulator drug combination elexacaftor/tezacaftor/ivacaftor (ELX/TEZ/IVA, ETI) has markedly improved clinical symptoms, but its broader molecular and systemic effects remain to be fully elucidated.

We employed mass spectrometry-based proteomics to compare the blood proteomes of CF patients treated with the earlier, less effective lumacaftor/ivacaftor (LUM/IVA) combination against those receiving the more potent ELX/TEZ/IVA therapy. Our analysis revealed both specific and common pharmacodynamic signatures associated with inflammation and metabolic processes under each treatment regimen. Notably, ELX/TEZ/IVA therapy exhibited more consistent alterations across patients that were directed towards profiles observed in healthy individuals.

Furthermore, by comparing sputum and blood proteomes of ELX/TEZ/IVA treated patients we identified counter-directional changes in the pulmonary surfactant-associated protein B, SFTPB, a potential biomarker of lung tissue repair, which also correlated with lung function improvements.

This study provides a comprehensive resource that enhances our understanding of CFTR modulator-driven proteome alterations, offering insights to both systemic and local protein regulation in CF. Our findings indicate that ELX/TEZ/IVA promotes broader systemic health improvements, providing critical insights that could shape future therapeutic strategies in CF.

## Introduction

Cystic fibrosis (CF) is a severe autosomal recessive disease that significantly impacts the life expectancy and overall well-being of affected individuals. The disease is caused mutations in the cystic fibrosis transmembrane conductance regulator (*CFTR*) gene. The CFTR gene encodes for a chloride and bicarbonate channel on the apical membrane of various epithelial tissues. The most common mutation, *F508del*, affects 85% of all CF patients ^1, 2^ and leads to disruptions in ion homeostasis, dehydration of epithelial surfaces, mucus thickening, chronic airway inflammation and recurrent infections ^2^. Together, these factors contribute to the challenges faced by individuals with CF. However, it is still unknown, if the systemic inflammation in CF is caused by CFTR dysfunction itself or a consequence of the overwhelming local inflammation in the lungs ^3–6^.

The most significant cause of disability and mortality in CF today is the mucus obstruction in the lungs, which is accompanied by a progressive deterioration of lung function ^3^. Beyond that, CF also exerts systemic effects by impairing the pancreatic, liver, and intestinal function, as well as affecting bone health ^2^.

In the last decade, CFTR modulators have been developed to pharmacological restore CFTR function ^1, 7^. Among these, lumacaftor/ivacaftor (LUM/IVA) and the more recent triple combination elexacaftor/tezacaftor/ivacaftor (ELX/TEZ/IVA) have shown promise. Lumacaftor, elexacaftor and tezacaftor act as CFTR correctors, enhancing the folding and trafficking of mutant CFTR, and are used in combination with ivacaftor, a potentiator of CFTR function.

The dual combination therapy LUM/IVA was approved in 2015 for patients with two copies of the *F508del* mutation (*F508del*/*F508del*), which constitutes approximately 50% of the CF population. This combination has shown modest improvements in ion homeostasis with reduced sweat chloride concentration by about 10-20% ^8–10^. Although a reduction in pulmonary exacerbations of about 30% was observed, there were only subtle effects on lung function (FEV_1_% predicted) ^11–13^. The triple combination therapy ELX/TEZ/IVA was approved recently for patients with at least one copy of the *F508del* mutation, making CFTR modulator therapy accessible to a majority of CF patients. The therapy has demonstrated significant positive outcomes, including a 40-50% reduction in sweat chloride concentration, a 10-20% improvement in FEV_1_% predicted, and a notable 60% decrease in pulmonary exacerbations ^14–16^. These benefits extend to various clinical parameters. Patients receiving ELX/TEZ/IVA therapy exhibit better lung ventilation and reduction in mucus plugging detected by magnetic resonance imaging (MRI) ^17–19^. Analysis of sputum from patients with CF treated with ELX/TEZ/IVA showed that these clinical findings are associated with positive changes in parameters such as rheology, microbiome, and proteome ^20–22^. These findings are also reflected in an increased life expectancy and quality of life ^5, 23, 24^.

In recent years, proteomics has emerged as a powerful tool for investigating complex disease mechanisms. Efficient protein extraction conditions employed in shotgun proteomics workflows and the untargeted detection of proteins by mass spectrometry make this technology in particular suitable for systems medicine studies ^25, 26^. Multiple groups Thus, have used proteomics has been frequently used to investigate the CF proteome in cell lines, primary cells, as well as blood and sputum samples ^20, 21, 27–31^. Several studies have investigated changes in the blood proteome longitudinally before and after pulmonary exacerbations (PEx), both with and without antibiotic treatment ^32–35^. These studies have highlighted the importance of inflammatory proteins, such as those involved in the complement pathway, C-reactive protein (CRP), alpha-1-antitrypsin (SERPINA1), and matrix metalloproteases ^36, 37, 35^ which have also been described in other biomarker studies alongside calprotectin (S100A8/9), serum amyloid A 1/2 protein (SAA1/2), and CD14 ^38–41^. Additionally, proteins related to lipid and vitamin transport and metabolism, such as apolipoproteins (apolipoprotein A-I (APOA1), apolipoprotein B-100 (APOB)), vitamin D-binding protein (GC), and retinol-binding protein 4 (RBP4), were shown to be altered in CF ^36, 42, 35, 39^. In the context of CF, blood proteomics is especially valuable as it provides an almost non-invasive means of assessing disease progression and treatment efficacy. This is particularly important as collecting sputum samples to understand local lung changes becomes increasingly challenging, especially in young children who may be unable to produce sputum.

The primary objective of this study was to identify and compare changes in the blood proteome composition resulting from LUM/IVA and ELX/TEZ/IVA therapies. These proteomics alterations were systematically correlated with the clinical efficacy of the treatments on key endpoints such as CFTR function (sweat chloride concentration) and lung function (FEV_1_% predicted). Further, the study aimed to differentiate between effects emerging from the lung, by including the corresponding sputum proteome from our previous research ^20^, and systemic effects observed in the blood. Our integrated analysis provides a valuable resource for understanding the pharmacodynamics of CFTR modulators, offering insights into the systemic and local protein regulation in CF (Fig. 1).

**Figure 1:**
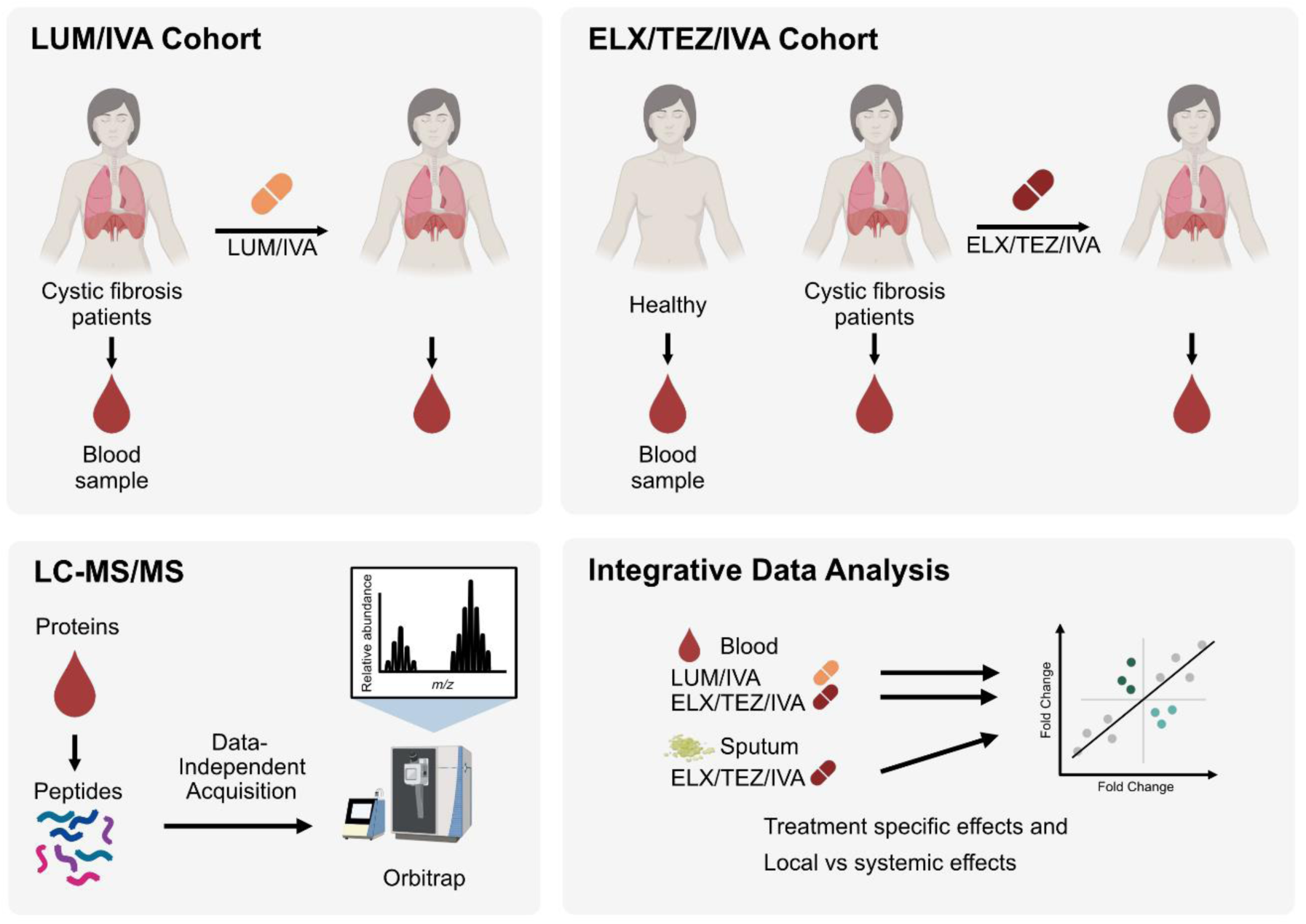
Study overview. Overview of the two patient cohorts treated with LUM/IVA (n = 32) or ELX/TEZ/IVA (n = 54) with proteomics workflow. The analysis included a comparison of both cohorts and an integrative analysis of overlapping patients from the ELX/TEZ/IVA cohort (n = 23), comparing blood data with sputum data from Schaupp et al. (2023) ^20^.

## Results

### Study Population Characteristics and Clinical Effects of LUM/IVA and ELX/TEZ/IVA

A total of 86 patients with CF and 11 healthy controls were analyzed in this study (Fig. 1). In the LUM/IVA cohort, all 32 patients were homozygous for the *F508del* mutation and were modulator-naïve at baseline. The ELX/TEZ/IVA cohort included 54 patients, of whom 55.6% were homozygous for the *F508del* mutation and 44.4% were heterozygous for the *F508del* mutation and a minimal function mutation. In this cohort, 53.7% were modulator-naïve, and 46.3% had received dual CFTR modulator therapy (LUM/IVA or TEZ/IVA) at baseline (Table 1). Consistent with previous studies ^8–13^, the LUM/IVA cohort did not exhibit a significant improvement in FEV_1_% predicted or a change in BMI following initiation of this therapy. In contrast, the ELX/TEZ/IVA cohort showed a significant increase in both parameters. Both cohorts exhibit a decrease in sweat chloride concentration with the ELX/TEZ/IVA group showing a more pronounced effect (Supplemental Fig. 1B). Notably, 23 patients who received ELX/TEZ/IVA were also included in a previously described sputum cohort ^20^.

**Table 1:**
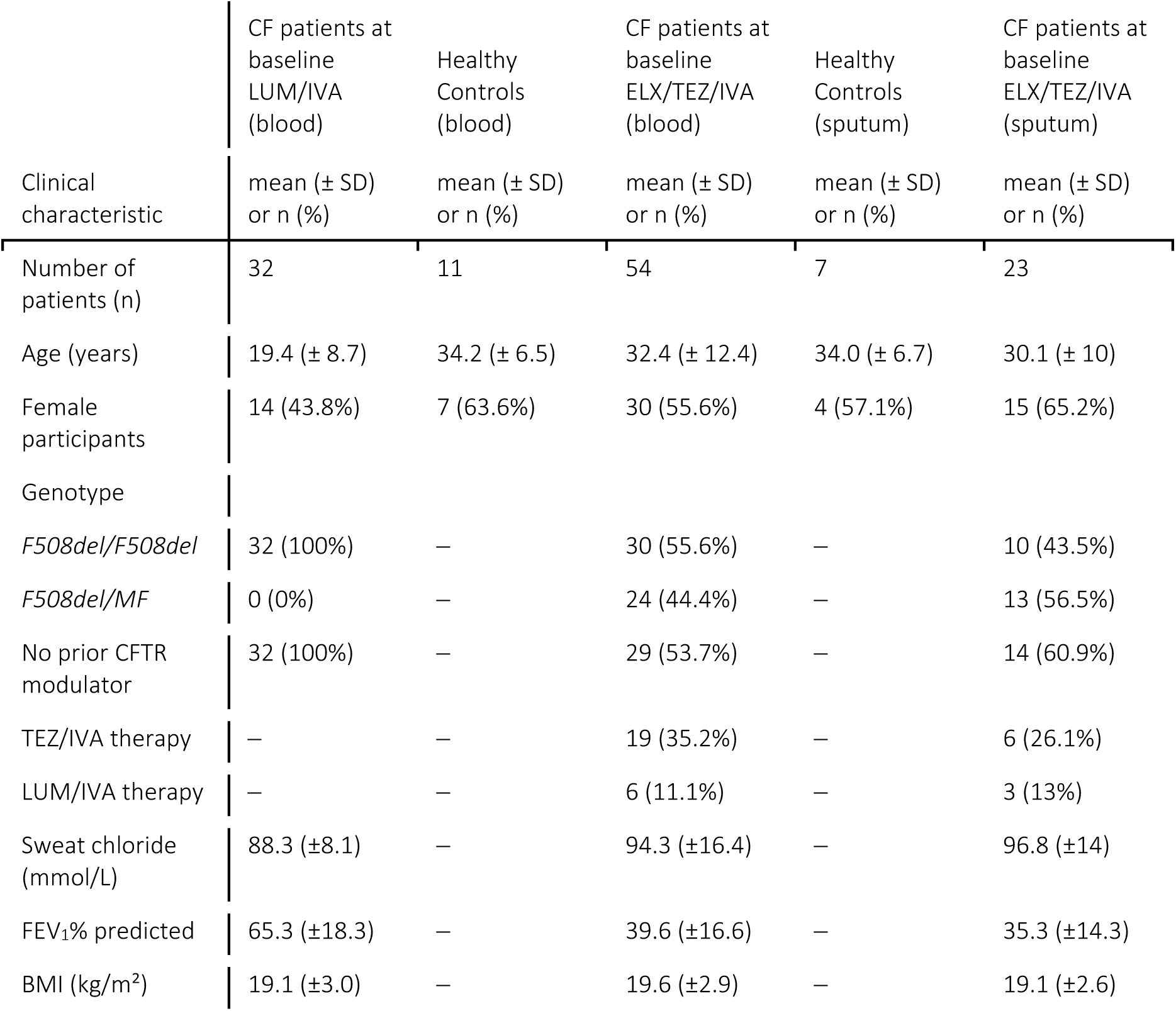
Baseline characteristics of the study population. Patients with cystic fibrosis (CF) in the ELX/TEZ/IVA sputum cohort are a subset of patients in the ELX/TEZ/IVA blood cohort.

### LUM/IVA Therapy Results in Moderate and Variable Changes in the Blood Proteome

To investigate the systemic effects of LUM/IVA therapy we performed a label-free shotgun mass-spectrometry analysis in data-independent acquisition mode (DIA) on plasma samples from 32 CF patients at baseline and 3 months after initiation of LUM/IVA therapy. Out of the 572 quantified proteins, 78 (13.6%) showed significant differences (Fig. 2A, Supplemental Table 1). These proteins were distributed over the entire intensity range (Supplemental Fig. 2C). Immunoglobulins were highly enriched in the group of downregulated proteins (Cluster 1), while hydrolases such as pantetheinase (VNN1) or mannan-binding lectin serine protease 2 (MASP2) were found among the upregulated proteins upon treatment (Cluster 2) (Fig. 2B).

**Figure 2:**
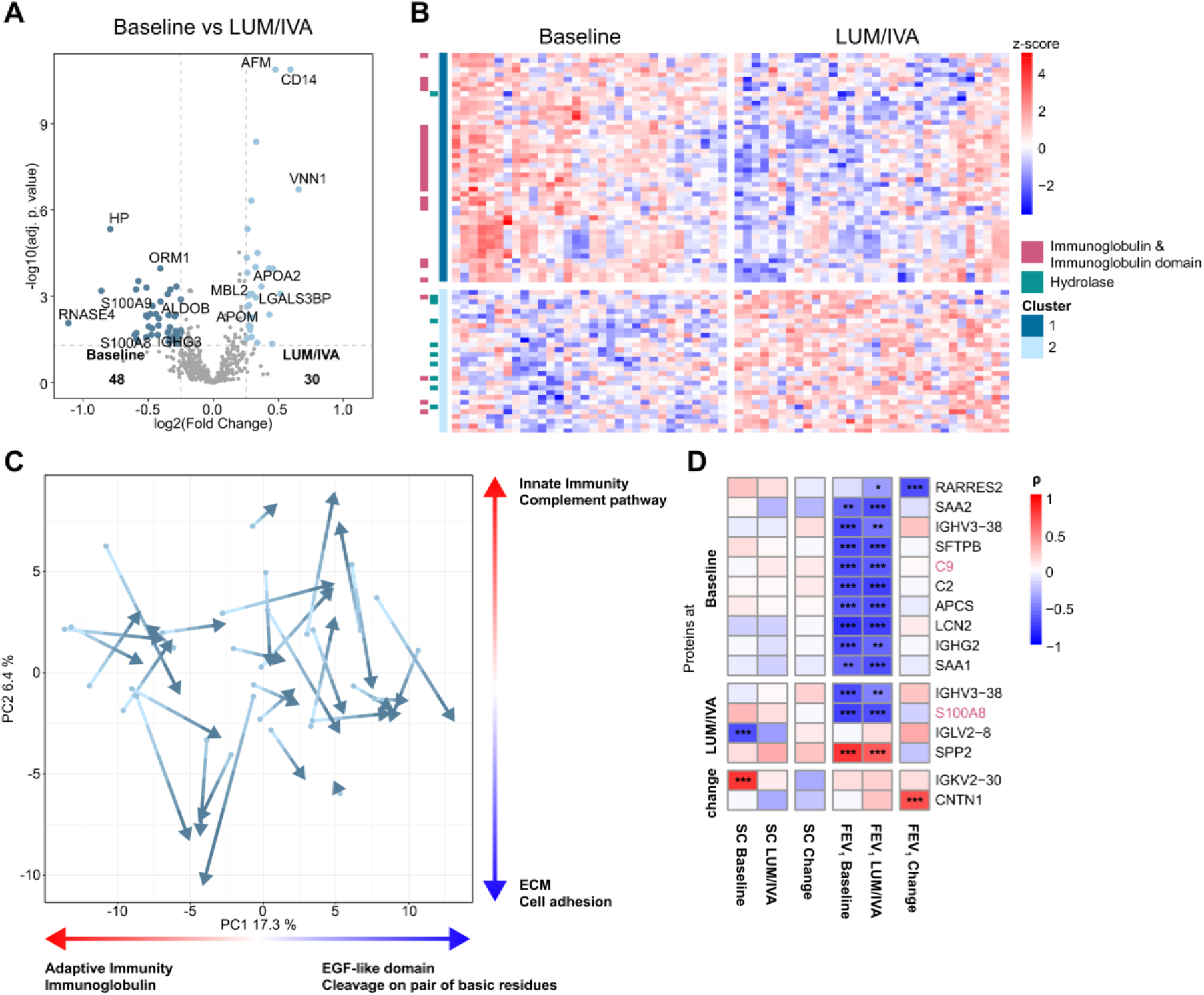
Moderate response in blood proteomes after 3 months of LUM/IVA treatment. (A) Volcano plot of proteins showing significantly increased or decreased (moderated t-test, Benjamini-Hochberg adjusted p < 0.05, ±0.25-log2(fold change)) protein abundances before and after 3 months of LUM/IVA treatment. (B) Heatmap of the 64 differential proteins (Fig. 2A). Hierarchical clustering was performed within the groups. Protein clusters are described by chosen UniProt terms enriched in one of the clusters. (C) Induced blood proteome changes of patients from baseline to treated visualized in principal component analysis. PCs are additionally described by 2 (top and bottom) of the most significantly enriched UniProt terms calculated by ssGSEA of eigenvectors. Arrows point from baseline to treated time points for the same patients. (D) Heatmap of spearman correlations between proteins at baseline, after treatment, or paired log2(fold changes) with sweat chloride concentration (SC) level or FEV_1_% predicted (FEV_1_) at baseline, after treatment or with the log2(fold changes) between baseline and treatment (change). Each row shows at least one significant correlation (p.adjust < 0.05). Protein names highlighted in red are proteins significantly different (p.adjust < 0.05) between Baseline and LUM/IVA. *** adj.p.val<0.001, ** adj.p.val<0.01, * adj.p.val< 0.05

Although significant shifts in the plasma proteome were observed from baseline to treatment in individual patients, there was no clear separation of the two groups on a global level, as shown by principal component analysis (PCA) (Fig. 2C). The first two principal components (PC1 and PC2) suggested a heterogeneous response, with only partial co-directional changes, particularly along PC1. Single-sample Gene Set Enrichment Analysis (ssGSEA) using UniProt terms of the eigenvectors from PC1 and PC2 showed that changes in immune responses, including adaptive and innate immunity, as well as changes in extracellular matrix (ECM) proteins (ECM, cell adhesion, EGF-like domain), contributed to the variance in PC1 and PC2 (Fig. 2C, Supplemental Table 2).

To relate the proteome changes the clinical characteristics, we performed correlation analysis of significantly changed proteins and the clinical parameters sweat chloride concentration, a biomarker of CFTR function, and FEV_1_% predicted, which serves as a marker of lung function. Two immunoglobulins (IGKV2.30 and IGLV2.8) correlated with sweat chloride levels at baseline. Multiple proteins correlated with lung function at baseline or after initiation of LUM/IVA, such as SFTPB and lipocalin-2 (LCN2), whereas contactin-1 (CNTN1) and retinoic acid receptor responder protein 2 (RARRES2) correlated with the change in FEV_1_% predicted (Fig. 2D).

### ELX/TEZ/IVA Induces Substantial Proteomic Shifts Towards a Healthier Blood Profile

To understand the systemic effect of ELX/TEZ/IVA therapy, we analyzed serum samples from 54 CF patients before and after 3 months of treatment and compared them to serum samples from 11 healthy individuals. Of the 559 quantified proteins, 196 (35.1%) exhibited significant differences between patients with CF at baseline and healthy individuals, while 68 proteins (12.2%) showed significant changes between baseline and 3 months after initiation of ELX/TEZ/IVA (Fig. 3A). Despite treatment, 101 proteins (18.1%) remained significantly different from healthy controls in the ELX/TEZ/IVA cohort. These changes spanned the entire intensity range (Supplemental Fig. 2B). 170 proteins (30.4%) were significantly different across the three groups (Fig. 3B). These proteins can be divided into two clusters. Cluster 1, representing proteins which were more abundant in CF patients at baseline in comparison to healthy individuals, was enriched in immunoglobulins and proteins involved in the complement pathway. On the other hand, cluster 2, which contained proteins less abundant in CF at baseline, was enriched in transport proteins such as transthyretin (TTR), GC, APOA1, and APOB.

**Figure 3:**
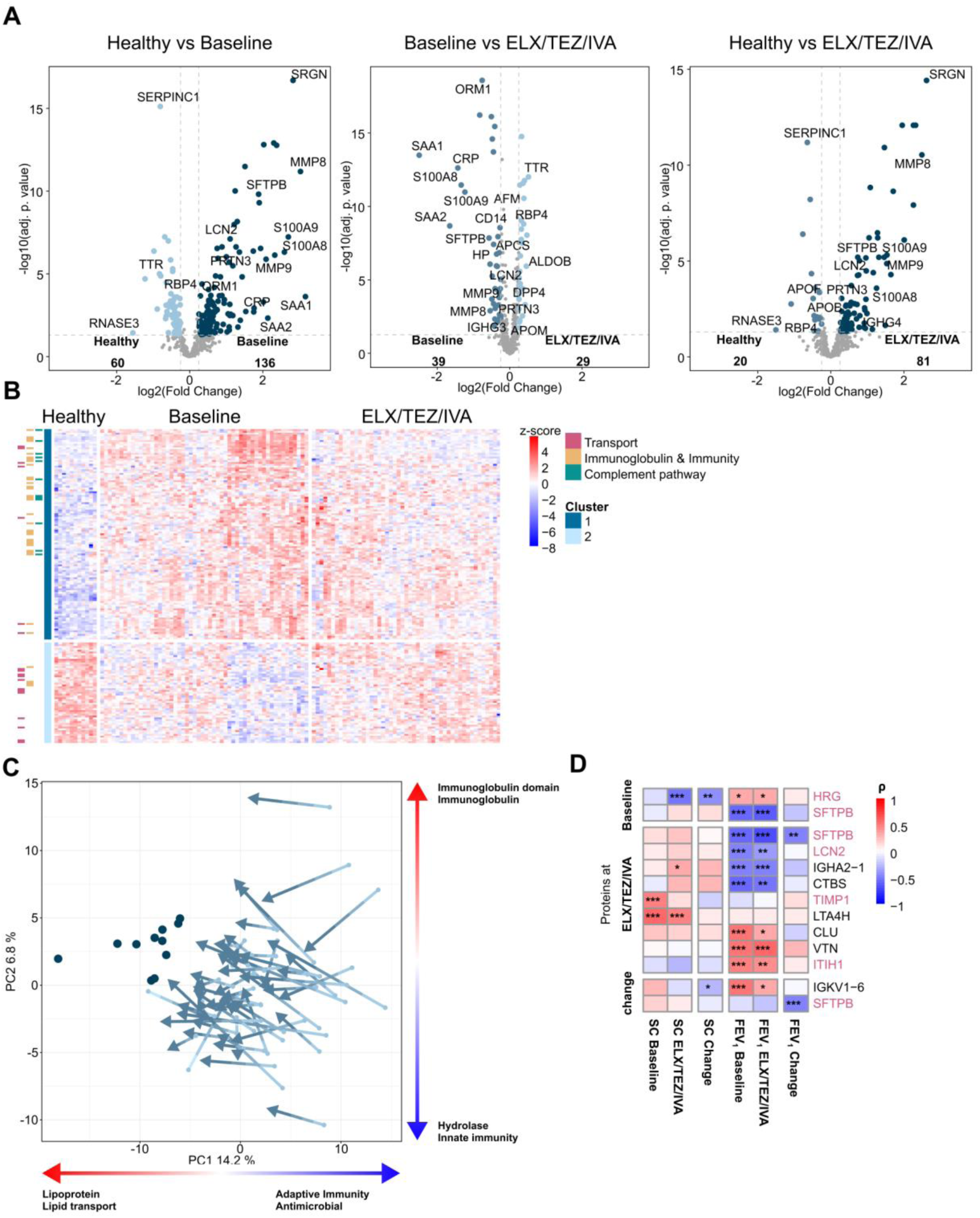
Strong and uniform response in blood proteomes after 3 months of ELX/TEZ/IVA treatment. (A) Volcano plots of proteins significantly increased or decreased (moderated t-test, Benjamini-Hochberg adjusted p < 0.05, ±0.25-log2(fold change)) protein abundances before treatment compared to healthy (left panel), before treatment compared to after 3 months of ELX/TEZ/IVA treatment (middle panel) and 3 months of ELX/TEZ/IVA treatment compared to healthy (right panel). (B) Heatmap of 170 significantly differential proteins between healthy, baseline and treated (moderated F-test, Benjamini-Hochberg adjusted p < 0.05). Hierarchical clustering was performed within the groups. Protein clusters are described by chosen UniProt terms enriched in one of the clusters. (C) Proteome regulation of patients from baseline to treated in comparison to healthy individuals visualized in principal component analysis. PCs are additionally described by 2 (top and bottom) of the most significantly enriched UniProt terms calculated by ssGSEA of eigenvectors. Arrows point from baseline to treated time points for the same patients. (D) Heatmap of spearman correlations between proteins at baseline, after treatment, paired log2(fold changes) with sweat chloride concentration (SC) level or FEV_1_% predicted (FEV_1_) at baseline, after treatment or with the log2(fold changes) between baseline and treatment (change). Each row shows at least one significant correlation (p.adjust < 0.05). Protein names highlighted in red are proteins significantly different (p.adjust < 0.05) between Baseline and ELX/TEZ/IVA. *** adj.p.val<0.001, ** adj.p.val<0.01, * adj.p.val< 0.05

Remarkably, most patient blood proteomes evolved upon treatment in the same direction, which was towards healthy blood proteomes (Fig. 3C). The primary contributors to the variance on PC1 were lipoproteins, proteins involved in lipid transport, as well as proteins involved in host immune defense.

Multiple proteins were found to correlate with sweat chloride concentration and FEV_1_% predicted (Fig. 3D). Specifically, histidine-rich glycoprotein (HRG) levels at baseline were significantly negatively correlated with sweat chloride levels in treated patients. Leukotriene-A4 hydrolase (LTA4H) levels under ELX/TEZ/IVA correlated with sweat chloride concentration at baseline and after 3 months of treatment, while metalloproteinase inhibitor 1 (TIMP1) levels in treated patients correlated with baseline sweat chloride concentration. Notably, all three proteins (HRG, LTA4H, and TIMP1) are involved in inflammatory processes ^43–45^.

Surfactant Protein B (SFTPB) exhibited negative correlations with FEV_1_% predicted both before and after 3 months of ELX/TEZ/IVA treatment. Furthermore, the levels of SFTPB in treated patients and the log2(fold change) of SFTPB also correlated with changes in FEV_1_% predicted. Additionally, the post-treatment levels of LCN2 correlated negatively, and inter-alpha-trypsin inhibitor heavy chain H1 (ITIH1) correlated positively with FEV_1_% predicted levels at baseline and after ELX/TEZ/IVA treatment.

### Stronger and Distinctive Response in ELX/TEZ/IVA than LUM/IVA Treated Patients

Since ELX/TEZ/IVA elicits a more robust response and improve patients’ clinical parameters and blood proteome, we sought to determine whether the differences were due to a larger effect size exceeding a critical threshold to induce a clinical response or if the response pattern was fundamentally distinct between both therapies.

Comparison of significantly different proteins between baseline and treated conditions in the two cohorts revealed that that the LUM/IVA cohort therapy led to a higher number of significantly different proteins (Fig. 4A), despite the larger effect size in ELX/TEZ/IVA treated patients (Fig. 4B). Interestingly, the difference between the two cohorts extended beyond the fold change. The majority of significantly different proteins did not overlap (Fig. 4), and some are proteins were regulated in the opposite direction (Fig. 4A, B).

**Figure 4:**
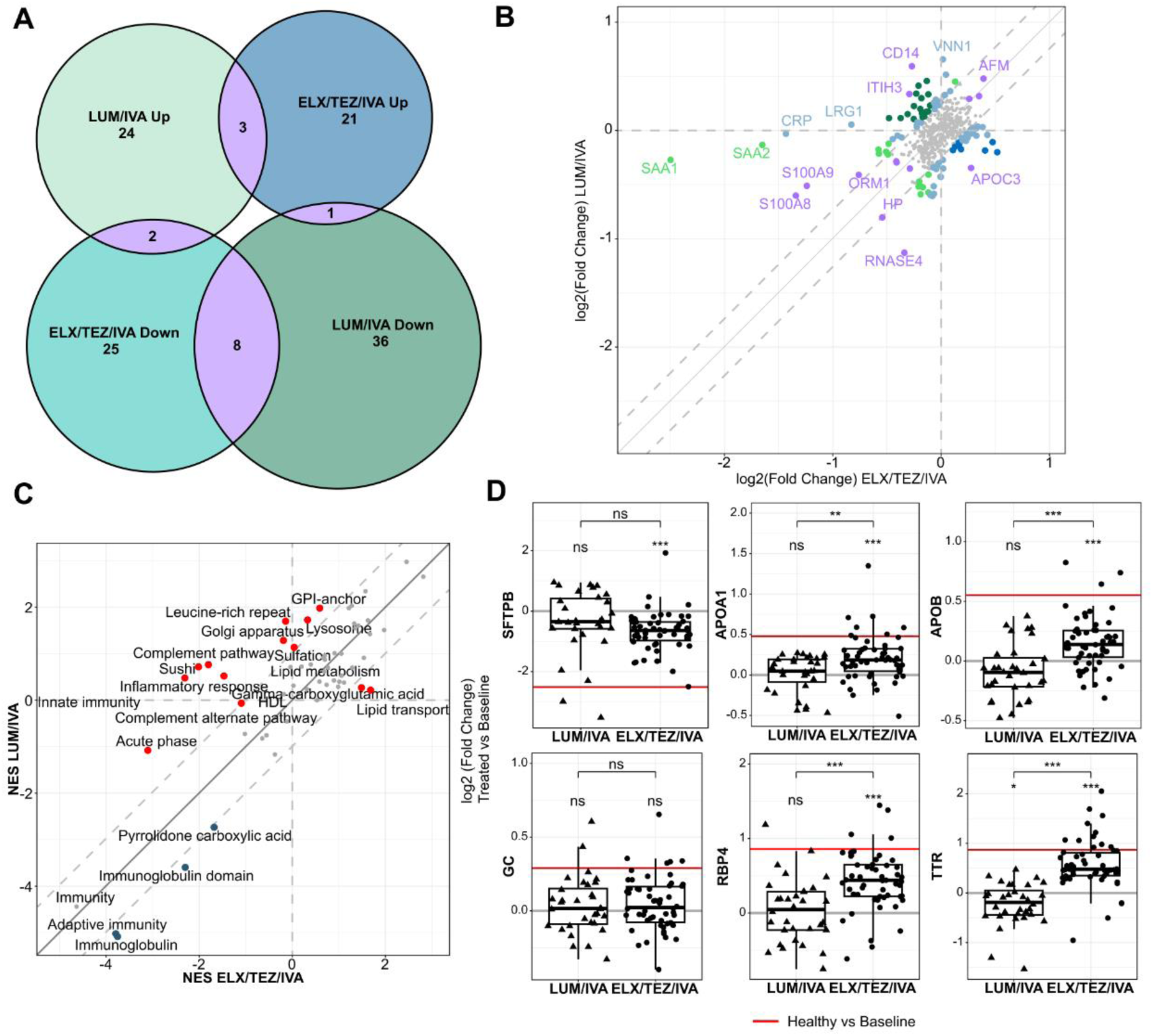
Innate immunity, lipid metabolism and vitamin transport are differentially affected by LUM/IVA and ELX/TEZ/IVA treatment. (A) Venn Diagram comparing proteins significantly up and down between 3 months of treatment and baseline in LUM/IVA and ELX/TEZ/IVA after correcting for differences in age. (B) Scatterplot of log2(fold changes) in commonly identified proteins between paired treated and baseline patients in the LUM/IVA or ELX/TEZ/IVA cohort after correcting for age as a confounder. Proteins highlighted have an absolute log2(fold change) difference of at least 0.25. Proteins changing in the same direction are highlighted in light green, proteins only changing in one sample (log2(fold change) in the other sample <0.1) are highlighted in light blue, proteins that changing counter directional are highlighted in dark green (higher in LUM/IVA) and dark blue (higher in ELX/TEZ/IVA). Significant proteins overlapping in the Venn diagram are highlighted in purple. (C) Scatterplot of normalized enrichment scores (NES) from ssGSEA analysis with UniProt keywords using paired log2(fold changes) from the LUM/IVA and the ELX/TEZ/IVA cohort. Terms significantly different between the LUM/IVA and the ELX/TEZ/IVA cohort are highlighted in red. Terms with a difference in NES below 1 are shown in grey. (D) Within and between cohort comparison of paired log2(fold changes) in LUM/IVA (triangles) and ELX/TEZ/IVA (dots) patients of representative proteins for different CF disease processes including the lung protein SFTPB, lipid metabolism related proteins (APOA1, APOB), and vitamin transport related proteins (GC, RPB4, TTR). Triangles and dots represent the paired log2(fold changes) between baseline and treated patients, red lines represent the log2(fold change) between CF baseline and healthy in the ELX/TEZ/IVA cohort. *** adj.p.val<0.001, ** adj.p.val<0.01, * adj.p.val< 0.05

Among the 471 proteins identified in both cohorts, 66 (14.0%) showed significant differences between the two groups (Supplemental Fig. 3E). While some proteins changed in the same direction (light green), many others showed opposite effects (dark green and dark blue) or only in one sample (light blue) (Fig. 4B). These differences were further reflected in the normalized enrichment scores (NES) of UniProt terms (Fig. 4C). Although most terms, such as ‘immunity’, ‘adaptive immunity’, ‘lipoproteins’, and ‘cell adhesion’, changed in the same direction in both therapies, many terms related to innate immunity, such as ‘complement pathway’, ‘complement alternate pathway’, and ‘inflammatory response’, were reduced after ELX/TEZ/IVA treatment but not after LUM/IVA treatment.

Next, we compared the changes of proteins representative for lung damage (SFTPB), lipid metabolism (APOA1, APOB), and vitamin transport (GC, RBP4, and TTR), which are key pathological pathways in CF, between the two treatment cohorts. Interestingly, none of the proteins, except for TTR, showed significant changes in the LUM/IVA cohort. Proteins related to lipid metabolism (APOA1, APOB) and vitamin transport (RBP4, and TTR), which were higher in healthy individuals compared to CF patients at baseline, exhibited significant increases with ELX/TEZ/IVA treatment but not with LUM/IVA (Fig. 4D).

### Differential Responses in Sputum and Blood Proteome after ELX/TEZ/IVA Treatment

For a subgroup of the ELX/TEZ/IVA cohort, previously published proteome data of matching sputum samples^20^ were available to determine the relationship between local treatment response in the lung with systemic changes. Additionally, we compared sputum and blood proteome data of the cohort including all patients (Supplemental Fig. 5).

Out of the 2541 proteins identified in the sputum samples and 558 proteins in the serum samples, 275 proteins overlapped (Supplemental Fig. 4A). Noteworthy, ELX/TEZ/IVA treatment induced more significant changes in protein abundance in serum than in sputum samples (Fig. 5A).

**Figure 5:**
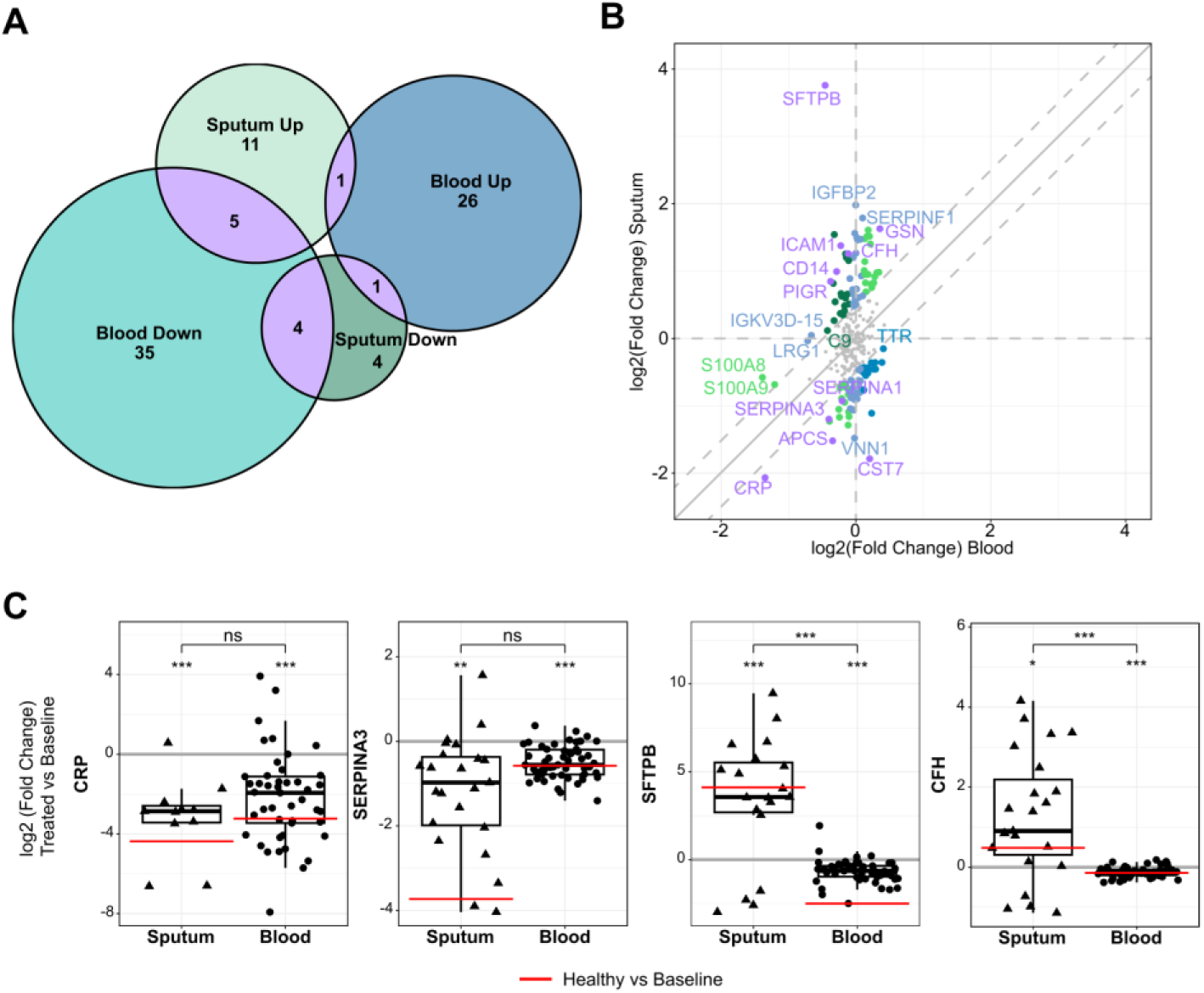
Differences in systemic (blood) and local (sputum) changes after treatment with ELX/TEZ/IVA. (A) Venn Diagram of overlapping proteins that are significantly up or down in the sputum and blood samples of ELX/TEZ/IVA treated patients (one sample moderated t-test, Benjamini-Hochberg adjusted p < 0.05). (B) Scatterplot of log2(fold changes) between treated and baseline patients in commonly identified proteins in plasma and sputum samples. Proteins highlighted have an absolute log2(fold change) difference of at least 0.25. Proteins changing in the same direction are highlighted in light green, proteins only changing in one sample (log2(fold change) in the other sample <0.1) are highlighted in light blue, proteins that changing counter directional are highlighted in dark green (Sputum) and dark blue (blood). Significant proteins overlapping in the Venn diagram are highlighted in purple. (C) Within and between cohort comparison of log2(fold changes) in sputum (triangles) and plasma (dots) of representative proteins developing in the same direction (inflammation marker (CRP, SERPINA3)) and opposite direction (lung protein SFTPB, and complement regulator CFH). Triangles and dots represent the paired log2(fold changes) between baseline and treated patients, red lines represent the log2(fold change) between baseline and healthy in the sputum or blood cohort. *** adj.p.val<0.001, ** adj.p.val<0.01, * adj.p.val< 0.05

Several inflammatory proteins, including CRP, serum amyloid P-component (APCS), SERPINA1, and alpha-1-antichymotrypsin (SERPINA3), showed significant differences and co-directional changes in both sputum and blood samples (Fig. 5A, B). In contrast, other proteins, including CD14, SFTPB, complement factor H (CFH), polymeric immunoglobulin receptor (PIGR), intercellular adhesion molecule 1 (ICAM1), and cystatin-F (CST7), displayed significant differences with opposite directionality (Fig. 5 A, B). Independent of the significance level, we compared the relation of effect size and found 89 proteins (32.4%) had a difference in absolute log2(fold change) of > 0.25 between the drug responses observed in sputum and blood datasets (highlighted proteins in Fig. 5B).

It is worth noting that while some inflammatory proteins, such as CRP and SERPINA3, were lower abundant in the sputum and blood of healthy individuals compared to CF patients at baseline and decreased upon ELX/TEZ/IVA treatment in both compartments. Others, such as CFH and SFTPB, were higher abundant in healthy sputum but lower abundant in the blood, and developed in opposite directions upon treatment (Fig. 5C). This development is particularly pronounced for SFTPB (Fig. 5B, C), which showed a significant correlation with lung function in the ELX/TEZ/IVA cohort (Fig. 5D). The peptide coverage of SFTPB is shown in Supplemental Fig. 6.These results illustrate the diverse impacts of ELX/TEZ/IVA therapy on local and systemic sites, as well as the dissimilarities in baseline protein levels in distinct tissues (Fig. 5C).

## Discussion

Here we applied a robust and highly scalable MS-based proteomics workflow to analyze blood samples from patients with CF who were either treated with CFTR modulator dual (LUM/IVA) or triple (ELX/TEZ/IVA) combination therapy. Previous CF research has primarily focused on analyzing specific blood proteins or conducting 2D-GE MS or MRM-MS, mostly comparing CF patients to healthy controls or PEx ^35, 36, 39, 46–48^. To our knowledge, this is the first study assessing the blood proteome in CF patients treated with LUM/IVA and ELX/TEZ/IVA.

We observed moderate effects in the LUM/IVA treated cohort (Fig. 2) and demonstrated that the proteome of patients in the ELX/TEZ/IVA cohort changes more uniformly its composition towards a healthier state upon treatment (Fig. 3). Notably, ELX/TEZ/IVA was more effective in modulating innate immune responses, leading to reduced airway and systemic inflammation (Fig. 4).

In addition, combining quantitative proteome analyses of blood and sputum samples allowed comparison of local (lung) and systemic changes upon treatment with ELX/TEZ/IVA (Fig. 5). Taken together, this integrated analysis provides a valuable resource for understanding the pharmacodynamics of different CFTR modulators and offer insights into the systemic and local protein regulation in CF.

## Effects of CFTR Modulator Therapy on Inflammation

A central challenge for CF patients is the persistent inflammation and infection of the lungs. Biomarkers such as CRP, S100A8, and S100A9 are widely discussed as indicators of CF severity. Particularly calprotectin (S100A8/A9) has been shown in multiple studies to be potentially useful for assessing the risk of pulmonary exacerbation (PEx) and monitoring disease progression. ^35, 39, 46, 47, 49–51^.

Despite the reduction in some immune related proteins with LUM/IVA, especially in immunoglobins, the overall impact on the proteome and clinical parameters remains limited. This is likely due to a sustained innate immunity response and a continuous activation of the complement pathway, alongside a lack of significant reduction in CRP levels.

However, our data shows decreased levels of the inflammation marker S100A8/9 after LUM/IVA and ETI treatment, which has been previously linked to a decreased risk of PEx ^52^, suggesting a protective function against further decline of lung function. In contrast, treatment with ELX/TEZ/IVA reduces proteins involved in both innate and adaptive immunity, which is also represented in serum proteome biomarkers such as CRP ^53^.

HRG, a protein involved in various inflammatory processes, is reduced in idiopathic pulmonary fibrosis ^54^, and sepsis ^43^ and has been shown to be capable of reducing systemic inflammation ^55, 56^. In the ELX/TEZ/IVA cohort, HRG levels were positively correlated with lung function and negatively correlated with sweat chloride concentration. HRG levels were increased in both cohorts after treatment, indicating a reduction in overall inflammation with both modulator treatments.

Another protein of interest is LTA4H, an inflammatory enzyme involved in the production of leukotriene B4 (LTB4) which in turn is part of neutrophil signaling ^44^. Although its levels did not change significantly with modulator treatment, LTA4H correlated with sweat chloride concentration in the ELX/TEZ/IVA cohort. LTA4H inhibitors have been tested in Phase 1 ^44^ and Phase 2 ^57^ trials, resulting in moderate improvements in the risk for PEx in patients with better lung function or modulator-treated CF patients. These moderate improvements may be due to its additional capacity to limit chronic neutrophilic airway inflammation ^58^.

The observed differences in inflammation markers between LUM/IVA and ELX/TEZ/IVA may be attributed to ongoing disease progression despite treatment in the LUM/IVA group. This could also account for the correlation between baseline inflammatory proteins and lung function (Fig. 2D).

### CFTR modulator therapy restores Protein levels Implicated in Malabsorption, and Vitamin Transport

CFTR dysfunction also affects mucus properties and clearance in the gastrointestinal tract and leads to abnormal pancreatic secretions in CF patients. This leads to malabsorption of nutrients due to decreased secretion of digestive enzymes from the pancreas in pancreatic insufficient patients ^59^.

Furthermore, inadequate synthesis of lipid processing proteins, such as APOA1 and APOB, in the intestinal tissue has been proposed as a contributing factor to insufficient lipid absorption. This may explain why pancreatic enzyme replacement therapy does not completely restore lipid absorption in CF patients ^60^.

Consistent with previous research ^36, 60^, we observed a reduction in APOA1 and APOB levels in CF patients compared to healthy controls, which was partially restored in patients treated with ELX/TEZ/IVA, but not in those treated with LUM/IVA. The significant difference in enrichment of the “lipid transport” UniProt term, including other apolipoproteins and phospholipid transfer protein (PLTP) further emphasizes the contrast between the two treatments.

In addition to fat resorption and dyslipidemia, transporters of fat-soluble vitamins, such as vitamin D-binding protein (GC), retinol-binding protein 4 (RBP4), and transthyretin (TTR), are reduced in blood of patients with CF ^36^. Although RBP4 is expressed in various tissues, studies suggested that circulating RBP4 is mostly derived from the liver and its level is highly regulated but dependent on liver retinol (vitamin A) levels ^61–63^. The levels of circulating retinol and RBP4 in complex with TTR depend on liver function and retinol absorption from food ^62, 64^. Interestingly, RBP4 has been proposed as a predictive marker for the severity of pulmonary exacerbations in chronic obstructive pulmonary disease (COPD) due to its association with the nutritional status of patients, affecting patient survival ^65^. In line with previous studies, RBP4 and TTR showed reduced abundance in CF at baseline compared to healthy ^36^ and increase upon treatment with ELX/TEZ/IVA but not LUM/IVA. The change in levels of RBP4 and TTR is most likely due to improved nutritional status after ELX/TEZ/IVA treatment, which is also reflected in increased lipoproteins and BMI after 3 months of treatment. However, we did not observe a significant increase in vitamin D-binding protein levels post-treatment, suggesting the need for continuous supplementation and regular monitoring of vitamin D levels in CF patients.

### Local vs Systemic Changes of Inflammation Markers upon ELX/TEZ/IVA treatment

The patients in this ELX/TEZ/IVA cohort partially overlap with those in a previously published cohort that described changes in sputum of CF patients after ELX/TEZ/IVA treatment ^20^. To distinguish between systemic effects and local changes in the lung, we compared the changes in overlapping patients and proteins. Proteins that were regulated in the same direction in both sputum and serum samples mainly belonged to inflammatory and immune pathways, such as CRP, APCS, and SERPINA3, AGT, AHSG, GSN, and SERPINF1. However, we observed that many inflammatory proteins exhibited changes that were different in magnitude and even in opposite direction between blood and sputum. This suggests that inflammation markers in the blood are at least partially independent of lung inflammation. Many of the blood inflammation markers observed here are not solely spill-over effects from the lung but indicative of the distinct systemic CF effects.

The protein CFH, which is known for inhibiting C3b, showed a reduction in serum post-treatment. In contrast to our results, CFH has been shown to be increased in lungs ^66^ and reduced in plasma ^67^ of patients with idiopathic pulmonary fibrosis. This reduction in CFH may be due to reduced systemic inflammation, also highlighted by the decrease in complement pathway enrichment in ELX/TEZ/IVA treated patients, which contrasts with its potential role in lung inflammation modulation. This comparison highlights that although lung inflammation may contribute to systemic inflammation and blood markers, many blood inflammation markers change independently of lung inflammation. This provides a potential explanation for the challenge in finding reproducible blood biomarkers for lung infection.

### Signs of lung repair

Pro-SFTPB has been discussed as a blood marker for lung damage, particularly in the context of lung cancer ^68–70^. One potential explanation for this association is the increased lung permeability resulting from various forms of lung injury ^71–73^. In COPD, SFTPB levels in sputum or bronchoalveolar lavage (BAL) samples correlate with lung function but do not show short-term changes upon treatment ^74^.

Interestingly, in our study, SFTPB was not only one of the two proteins (SFTPB, LCN2) that correlated in multiple lung function comparisons in both blood cohorts, and especially also with the change in lung function in the ELX/TEZ/IVA cohort. Additionally, it was one of the proteins that changed in opposite direction when comparing changes in the blood and sputum proteome.

Within the ELX/TEZ/IVA cohort, decreased SFTPB levels in serum alongside increased levels in sputum, coupled with an improvement in lung function, could represent a potential indication of reduced lung permeability and, consequently, lung repair. This phenomenon was not observed in the LUM/IVA cohort. These findings suggests that SFTPB in blood samples may serve as a marker for evaluating lung damage and potentially repair, warranting further validation in future studies.

This study demonstrates that the superior efficacy of ELX/TEZ/IVA over LUM/IVA treatment is clearly reflected in the blood proteome, even though 46% of the patients in the ELX/TEZ/IVA cohort received modulator treatment at baseline. The findings highlighted that ELX/TEZ/IVA resulted in a greater improvement in blood inflammation markers than LUM/IVA. Moreover, the study showed that local changes in the lung only partially aligned with systemic changes, emphasizing the challenge of identifying suitable blood biomarkers for lung diseases. In addition, the study provides a comprehensive proteomic analysis of longitudinally collected blood samples from treated CF patients, offering a valuable reference for future research. It also serves as a resource for understanding how effective treatment may influence patient health, potentially reducing the risk of severe lung damage over time. However, the study also underscores the need for continued development of therapies, as significant differences remain between the blood proteomes of treated CF patients and healthy individuals, particularly in inflammatory proteins and potential lung injury markers such as SFTPB.

## Resource Availability

### Lead contact

Requests for further information and resources should be directed to the lead contact, Philipp Mertins

### Materials Availibility

This study did not generate new unique reagents.

### Data and Code Availability

The mass spectrometry proteomics data have been deposited to the ProteomeXchange Consortium via the PRIDE ^75^ partner repository. Data are available via ProteomeXchange with identifier PXD056648.

## Acknowledgments

The authors thank the patients with CF for their participation in this study; M. Daniltchenko, M. Drescher, K. Seidel and J. Tattersall-Wong (Charité –Universitätsmedizin Berlin, Berlin, Germany) for excellent technical assistance; C. Labitzke and E. Halver (Charité – Universitätsmedizin Berlin, Berlin, Germany) for excellent documentation. Schematic Figure was created with BioRender.com.

## Author Contributions

Conceptualization: K.F., M.K., M.Z., S.Y.G., M.A.M. and P.M.

Formal analysis: K.F., M.K., M.Z. and O.P.

Data curation: K.F., M.K., S.N., S.Y.G.

Investigation: K.F., M.K., M.Z., O.P., S.N., J.D., L.S., J.R., S.T., S.H., O.S, M.S, S.Y.G., M.A.M.

Resources: J.D., L.S., J.R., S.T., S.H., O.S, M.S, S.Y.G., M.A.M., P.M.

Supervision: S.Y.G, P.M., M.A.M.

Visualization: K.F.

Writing – original draft: K.F., S.Y.G, M.A.M., P.M.

Writing – review & editing: all authors

## Methods

### Study design and participants

The patients included in this study are part of two prospective longitudinal observational studies in patients with CF followed at the CF Centers at University Hospital Heidelberg, Germany (LUM/IVA cohort, ^9^) and at Charité - Universitätsmedizin Berlin, Germany (ELX/TEZ/IVA cohort, ^76^). The studies were approved by the ethics committee of the University of Heidelberg (S-370/2011) and the Charité - Universitätsmedizin Berlin (EA2/220/18). Written informed consent was obtained from all patients, their parents or guardians. Patients were eligible to take part in the LUM/IVA study, if they were homozygous for the *F508del* mutation, had no prior exposure to LUM/IVA and were willing to remain on a stable medication regimen and administration of LUM/IVA according to the EMA-approved patient label. In the ELX/TEZ/IVA study, patients were eligible, if they were compound-heterozygous for *F508del* and a minimal function mutation or homozygous for *F508del*, had no prior exposure to ELX/TEZ/IVA and were willing to remain on a stable medication regimen including ELX/TEZ/IVA according to the patient labeling and the prescribing information for the duration of study participation. Exclusion criteria for both studies were an acute respiratory infection or pulmonary exacerbation at baseline. In both studies, blood samples were collected at baseline and 3 months after initiation of therapy. Due to limited amounts of samples, the cohorts were measured in different types of blood derived samples (serum and plasma). Additionally, sweat chloride concentration, lung function and BMI were assessed at baseline and after 3 months of either LUM/IVA or ELX/TEZ/IVA treatment. Healthy control subjects were age- and sex-matched non-smoking volunteers.

### Proteomic measurements

#### Sample preparation

Serum and plasma samples were processed separately. Samples were diluted 1:10 in HPLC grade water for further processing. 10 µl of the dilution were denatured in SDC buffer (final concentration: 1% SDC, 10 mM DTT, 40 mM CAA, 100 mM Tris (pH 8)) by heating to 95°C for 10 minutes. After cooling the samples LysC and Trypsin were added with a peptide to enzyme ratio of 50:1 and samples digested overnight at 37°C. The digest was stopped by adding formic acid (FA) to a final concentration of 1%. The samples were diluted in Buffer A (3% Acetonitrile, 0.1% FA) and centrifuged for 10 minutes at 12 000 rpm. The peptide containing supernatant was collected and cleaned up using stage tips ^77^.

#### DIA analyses

Peptides were eluted from stage tips using 80 µl Buffer B (90% Acetonitrile, 0.1% FA), dried, resuspended in 10 µl Buffer A. 2 µl of each sample were injected. Peptides were separated on an 33 min gradient with increasing acetonitrile concentration using an EASY-nLC 1200 System coupled to an Orbitrap HF-X mass spectrometer (Thermo Fisher Scientific, Waltham, MA, USA) running on data-independent acquisition mode (DIA) as described with some adjustments ^78^. Briefly, one MS1 scan at 120k resolution with an AGC target of 3×10^6^ and max. injection time of 60 ms in the range of 350 to 1650 m/z was acquired, followed by 40 DIA scans with variable segment widths adjusted to the precursor density. The scan resolution in the Orbitrap was set to 30k with an AGC target of 3×10^6^ and max. injection time of 35 ms. The stepped HCD collision energy was set to 25.5,27 and 30%.

#### Plasma library preparation

For each cohort a separate peptide library was measured. 5 µl of each diluted sample within a cohort were pooled (2 mg in total) and digested as described above, cleaned up SepPak C18 columns, and fractionated using high-pH reversed phase off-line chromatography (1290 Infinity, Agilent; XBridge C18 4.6 mm × 250 mm column [Waters, 3.5 µm bead size]) into 196 fractions, which were concatenate into 52 analytical fractions. The fractions were dried and resuspended in Buffer A.

#### Depletion of plasma library

In order to increase library coverage, immunoaffinity depletion of 14 most abundant proteins followed by the next ∼50 moderately abundant proteins using IgY14 LC20 and Supermix LC10 columns (Sigma-Aldrich, St.Louis, MO) was performed as described ^79^. Briefly, six rounds of tandem depletion of 400 µl patient plasma as well as commercial human plasma samples were performed on an Agilent 1260 HPLC (Agilent, Santa Clara, CA) system using Dilution, Stripping, and Neutralization buffers provided by the manufacturer and following manufacturer’s instructions (Sigma-Aldrich). Flow throughs of the Supermix column from each round representing depleted plasma were combined, concentrated and buffer exchanged to 50 mm Ammonium Bicarbonate to the original volume (400 μl) using Amicon 3K concentrators (Millipore, Billerica, MA). The final protein sample was digested and fractionated as described above.

#### DDA analyses of library samples

One µg peptide of depleted and non-depleted library peptide fractions was measured on an Orbitrap HF-X mass spectrometer (Thermo Fisher Scientific, Waltham, MA, USA) running on data-dependent acquisition mode, using the same LC parameters as described for DIA measurements. The MS parameters were the following: MS1 scan at 60K resolution with AGC target of 3×10^6^ and maximum injection time of 10 ms, followed by twenty MS2 scan at 15K resolution with AGC target of 1×10^5^ and maximum injection time of 22 ms. Dynamic exclusion was set to 20 seconds.

#### RAW Data analyses

A combined spectral library with the two cohorts and a depleted library were created using Spectronaut (version 18.2) using the UniProt database including isoforms (2022-03) and a universal protein contaminants list. Modifications were set to Carbamidomethyl (C) as fixed and Acetyl (Protein N-term), Deamidation (NQ), and Oxidation (M) as variable modification. For the analysis of DIA data, precursor filtering was done using the *Q* value setting. Proteins were exported with and without background imputation, log2-transformed and data was filtered to contain at least 50% valid values in the non-imputed data frame. Remaining missing values were imputed using down-shift imputation by random draw from the Gaussian distribution with 0.3x standard deviation and downshift of 1.8x standard deviation of the observed values per sample ^80^. MaxLFQ was used for quantification.

DDA data from sputum samples were analyzed as described before ^20^. In summary, the MaxQuant software package (Ver. 2.0.3.0; Max Planck Institute of Biochemistry, Martinsried, Germany) ^81^ was used to analyze raw data from included patient samples. A UniProt database (2023-03) was included in the search. Modifications were defined as follows: oxidation (M), N-terminal acetylation, and deamidation (N,Q) as variable modifications, and carbamidomethyl cysteine as a fixed modification. Proteins with only one peptide were allowed. The search employed the “Match between runs” and label-free quantitation (LFQ) algorithm.

Further downstream analysis was performed in R (V 4.2.2.). MaxQuant output was filtered for “Reverse”, “Potential Contaminant”, and “Only identified by site”. Proteins identified with only one peptide were filtered for at least 5 identifications by MS/MS and a minimum andromeda score of 20. In the Spectronaut output contaminants were removed. For significance calling moderated t-test (limma package ^82^) was applied. Benjamini-Hochberg method was used to adjust p-values for multiple comparisons. For the different comparisons the cut-offs are described in the figure legend. For single-sample gene-set enrichment analysis (ssGSEA ^83^) a curated gmt file with UniProt keywords (2021-04) was used. Because of the differences in sample collection between the two cohorts and the differences in age we used a linear model to adjust for age and compared log2(fold changes) of proteins identified in both cohorts.

#### Comparison of serum and plasma samples

Matching serum and plasma samples from three baseline CF patient samples were digested as described above and measured on an Orbitrap HF-X mass spectrometer (Thermo Fisher Scientific, Waltham, MA, USA) running on data-dependent acquisition mode, using the same parameters as described for library sample analyses. Raw data of patient and plasma library fractions were processed using MaxQuant software package (v1.6.0.1) and a human UniProt database (2019) was used for the search. The search included variable modifications of oxidation (M), N-terminal acetylation, deamidation (N and Q) and fixed modification of carbamidomethyl cysteine. The FDR was set to 1% for peptide and protein identifications. Unique and razor peptides were considered for quantification. MS2 identifications were transferred from library runs to patient samples with the “match between runs” option. The integrated LFQ (label-free) quantitation algorithm was applied. Reverse hits, contaminants and proteins only identified by site were filtered out. Downstream analysis was done as described above.

## References

1. Boeck, K. de, Zolin, A., Cuppens, H., Olesen, H.V., and Viviani, L. (2014). The relative frequency of CFTR mutation classes in European patients with cystic fibrosis. Journal of cystic fibrosis : official journal of the European Cystic Fibrosis Society 13, 403–409. 10.1016/j.jcf.2013.12.003.

2. Mall, M.A., Burgel, P.-R., Castellani, C., Davies, J.C., Salathe, M., and Taylor-Cousar, J.L. (2024). Cystic fibrosis. Nature reviews. Disease primers 10, 53. 10.1038/s41572-024-00538-6.

3. Bacalhau, M., Camargo, M., Magalhães-Ghiotto, G.A.V., Drumond, S., Castelletti, C.H.M., and Lopes-Pacheco, M. (2023). Elexacaftor-Tezacaftor-Ivacaftor: A Life-Changing Triple Combination of CFTR Modulator Drugs for Cystic Fibrosis. Pharmaceuticals (Basel, Switzerland) 16. 10.3390/ph16030410.

4. Grasemann, H., and Ratjen, F. (2023). Cystic Fibrosis. The New England journal of medicine 389, 1693–1707. 10.1056/NEJMra2216474.

5. Bell, S.C., Mall, M.A., Gutierrez, H., Macek, M., Madge, S., Davies, J.C., Burgel, P.-R., Tullis, E., Castaños, C., and Castellani, C., et al. (2020). The future of cystic fibrosis care: a global perspective. The Lancet. Respiratory medicine 8, 65–124. 10.1016/S2213-2600(19)30337-6.

6. Ratjen, F., Saiman, L., Mayer-Hamblett, N., Lands, L.C., Kloster, M., Thompson, V., Emmett, P., Marshall, B., Accurso, F., and Sagel, S., et al. (2012). Effect of azithromycin on systemic markers of inflammation in patients with cystic fibrosis uninfected with Pseudomonas aeruginosa. Chest 142, 1259–1266. 10.1378/chest.12-0628.

7. Boeck, K. de, Munck, A., Walker, S., Faro, A., Hiatt, P., Gilmartin, G., and Higgins, M. (2014). Efficacy and safety of ivacaftor in patients with cystic fibrosis and a non-G551D gating mutation. Journal of cystic fibrosis : official journal of the European Cystic Fibrosis Society 13, 674–680. 10.1016/j.jcf.2014.09.005.

8. Sagel, S.D., Khan, U., Heltshe, S.L., Clancy, J.P., Borowitz, D., Gelfond, D., Donaldson, S.H., Moran, A., Ratjen, F., and VanDalfsen, J.M., et al. (2021). Clinical Effectiveness of Lumacaftor/Ivacaftor in Patients with Cystic Fibrosis Homozygous for F508del-CFTR. A Clinical Trial. Annals of the American Thoracic Society 18, 75–83. 10.1513/AnnalsATS.202002-144OC.

9. Graeber, S.Y., Dopfer, C., Naehrlich, L., Gyulumyan, L., Scheuermann, H., Hirtz, S., Wege, S., Mairbäurl, H., Dorda, M., and Hyde, R., et al. (2018). Effects of Lumacaftor-Ivacaftor Therapy on Cystic Fibrosis Transmembrane Conductance Regulator Function in Phe508del Homozygous Patients with Cystic Fibrosis. American journal of respiratory and critical care medicine 197, 1433–1442. 10.1164/rccm.201710-1983OC.

10. Clancy, J.P., Rowe, S.M., Accurso, F.J., Aitken, M.L., Amin, R.S., Ashlock, M.A., Ballmann, M., Boyle, M.P., Bronsveld, I., and Campbell, P.W., et al. (2012). Results of a phase IIa study of VX-809, an investigational CFTR corrector compound, in subjects with cystic fibrosis homozygous for the F508del-CFTR mutation. Thorax 67, 12–18. 10.1136/thoraxjnl-2011-200393.

11. Wainwright, C.E., Elborn, J.S., Ramsey, B.W., Marigowda, G., Huang, X., Cipolli, M., Colombo, C., Davies, J.C., Boeck, K. de, and Flume, P.A., et al. (2015). Lumacaftor-Ivacaftor in Patients with Cystic Fibrosis Homozygous for Phe508del CFTR. The New England journal of medicine 373, 220–231. 10.1056/NEJMoa1409547.

12. Konstan, M.W., McKone, E.F., Moss, R.B., Marigowda, G., Tian, S., Waltz, D., Huang, X., Lubarsky, B., Rubin, J., and Millar, S.J., et al. (2017). Assessment of safety and efficacy of long-term treatment with combination lumacaftor and ivacaftor therapy in patients with cystic fibrosis homozygous for the F508del-CFTR mutation (PROGRESS): a phase 3, extension study. The Lancet. Respiratory medicine 5, 107–118. 10.1016/S2213-2600(16)30427-1.

13. Rayment, J.H., Asfour, F., Rosenfeld, M., Higgins, M., Liu, L., Mascia, M., Paz-Diaz, H., Tian, S., Zahigian, R., and McColley, S.A. (2022). A Phase 3, Open-Label Study of Lumacaftor/Ivacaftor in Children 1 to Less Than 2 Years of Age with Cystic Fibrosis Homozygous for F508del-CFTR. American journal of respiratory and critical care medicine 206, 1239–1247. 10.1164/rccm.202204-0734OC.

14. Middleton, P.G., Mall, M.A., Dřevínek, P., Lands, L.C., McKone, E.F., Polineni, D., Ramsey, B.W., Taylor-Cousar, J.L., Tullis, E., and Vermeulen, F., et al. (2019). Elexacaftor-Tezacaftor-Ivacaftor for Cystic Fibrosis with a Single Phe508del Allele. The New England journal of medicine 381, 1809– 1819. 10.1056/NEJMoa1908639.

15. Heijerman, H.G.M., McKone, E.F., Downey, D.G., van Braeckel, E., Rowe, S.M., Tullis, E., Mall, M.A., Welter, J.J., Ramsey, B.W., and McKee, C.M., et al. (2019). Efficacy and safety of the elexacaftor plus tezacaftor plus ivacaftor combination regimen in people with cystic fibrosis homozygous for the F508del mutation: a double-blind, randomised, phase 3 trial. The Lancet 394, 1940–1948. 10.1016/S0140-6736(19)32597-8.

16. Griese, M., Costa, S., Linnemann, R.W., Mall, M.A., McKone, E.F., Polineni, D., Quon, B.S., Ringshausen, F.C., Taylor-Cousar, J.L., and Withers, N.J., et al. (2021). Safety and Efficacy of Elexacaftor/Tezacaftor/Ivacaftor for 24 Weeks or Longer in People with Cystic Fibrosis and One or More F508del Alleles: Interim Results of an Open-Label Phase 3 Clinical Trial. American journal of respiratory and critical care medicine 203, 381–385. 10.1164/rccm.202008-3176LE.

17. Graeber, S.Y., Renz, D.M., Stahl, M., Pallenberg, S.T., Sommerburg, O., Naehrlich, L., Berges, J., Dohna, M., Ringshausen, F.C., and Doellinger, F., et al. (2022). Effects of Elexacaftor/Tezacaftor/Ivacaftor Therapy on Lung Clearance Index and Magnetic Resonance Imaging in Patients with Cystic Fibrosis and One or Two F508del Alleles. American journal of respiratory and critical care medicine 206, 311–320. 10.1164/rccm.202201-0219OC.

18. Streibel, C., Willers, C.C., Pusterla, O., Bauman, G., Stranzinger, E., Brabandt, B., Bieri, O., Curdy, M., Bullo, M., and Frauchiger, B.S., et al. (2023). Effects of elexacaftor/tezacaftor/ivacaftor therapy in children with cystic fibrosis - a comprehensive assessment using lung clearance index, spirometry, and functional and structural lung MRI. Journal of cystic fibrosis : official journal of the European Cystic Fibrosis Society 22, 615–622. 10.1016/j.jcf.2022.12.012.

19. Stahl, M., Dohna, M., Graeber, S.Y., Sommerburg, O., Renz, D.M., Pallenberg, S.T., Voskrebenzev, A., Schütz, K., Hansen, G., and Doellinger, F., et al. (2024). Impact of elexacaftor/tezacaftor/ivacaftor therapy on lung clearance index and magnetic resonance imaging in children with cystic fibrosis and one or two F508del alleles. European Respiratory Journal 64. 10.1183/13993003.00004-2024.

20. Schaupp, L., Addante, A., Völler, M., Fentker, K., Kuppe, A., Bardua, M., Duerr, J., Piehler, L., Röhmel, J., and Thee, S., et al. (2023). Longitudinal Effects of Elexacaftor/Tezacaftor/Ivacaftor on Sputum Viscoelastic Properties, Airway Infection and Inflammation in Patients with Cystic Fibrosis. European Respiratory Journal. 10.1183/13993003.02153-2022.

21. Maher, R.E., Barry, P.J., Emmott, E., Jones, A.M., Lin, L., McNamara, P.S., Smith, J.A., and Lord, R.W. (2023). Influence of highly effective modulator therapy on the sputum proteome in cystic fibrosis. Journal of cystic fibrosis : official journal of the European Cystic Fibrosis Society. 10.1016/j.jcf.2023.10.019.

22. Casey, M., Gabillard-Lefort, C., McElvaney, O.F., McElvaney, O.J., Carroll, T., Heeney, R.C., Gunaratnam, C., Reeves, E.P., Murphy, M.P., and McElvaney, N.G. (2023). Effect of elexacaftor/tezacaftor/ivacaftor on airway and systemic inflammation in cystic fibrosis. Thorax 78, 835–839. 10.1136/thorax-2022-219943.

23. Piehler, L., Thalemann, R., Lehmann, C., Thee, S., Röhmel, J., Syunyaeva, Z., Stahl, M., Mall, M.A., and Graeber, S.Y. (2023). Effects of elexacaftor/tezacaftor/ivacaftor therapy on mental health of patients with cystic fibrosis. Frontiers in pharmacology 14, 1179208. 10.3389/fphar.2023.1179208.

24. Graeber, S.Y., and Mall, M.A. (2023). The future of cystic fibrosis treatment: from disease mechanisms to novel therapeutic approaches. The Lancet 402, 1185–1198. 10.1016/S0140-6736(23)01608-2.

25. Geyer, P.E., Kulak, N.A., Pichler, G., Holdt, L.M., Teupser, D., and Mann, M. (2016). Plasma Proteome Profiling to Assess Human Health and Disease. Cell systems 2, 185–195. 10.1016/j.cels.2016.02.015.

26. Messner, C.B., Demichev, V., Wendisch, D., Michalick, L., White, M., Freiwald, A., Textoris-Taube, K., Vernardis, S.I., Egger, A.-S., and Kreidl, M., et al. (2020). Ultra-High-Throughput Clinical Proteomics Reveals Classifiers of COVID-19 Infection. Cell systems 11, 11–24.e4. 10.1016/j.cels.2020.05.012.

27. Hoppe, J.E., Wagner, B.D., Kirk Harris, J., Rowe, S.M., Heltshe, S.L., DeBoer, E.M., and Sagel, S.D. (2022). Effects of ivacaftor on systemic inflammation and the plasma proteome in people with CF and G551D. Journal of cystic fibrosis : official journal of the European Cystic Fibrosis Society 21, 950–958. 10.1016/j.jcf.2022.03.012.

28. Braccia, C., Christopher, J.A., Crook, O.M., Breckels, L.M., Queiroz, R.M.L., Liessi, N., Tomati, V., Capurro, V., Bandiera, T., and Baldassari, S., et al. (2022). CFTR Rescue by Lumacaftor (VX-809) Induces an Extensive Reorganization of Mitochondria in the Cystic Fibrosis Bronchial Epithelium. Cells 11. 10.3390/cells11121938.

29. Trappe, A., Lakkappa, N., Carter, S., Dillon, E., Wynne, K., McKone, E., McNally, P., and Coppinger, J.A. (2023). Investigating serum extracellular vesicles in Cystic Fibrosis. Journal of cystic fibrosis : official journal of the European Cystic Fibrosis Society 22, 674–679. 10.1016/j.jcf.2023.02.005.

30. Pedrazzi, M., Vercellone, S., Barberis, E., Capraro, M., Tullio, R. de, Cresta, F., Casciaro, R., Castellani, C., Patrone, M., and Marengo, E., et al. (2021). Identification of Potential Leukocyte Biomarkers Related to Drug Recovery of CFTR: Clinical Applications in Cystic Fibrosis. International Journal of Molecular Sciences 22. 10.3390/ijms22083928.

31. Hisert, K.B., Birkland, T.P., Schoenfelt, K.Q., Long, M.E., Grogan, B., Carter, S., Liles, W.C., McKone, E.F., Becker, L., and Manicone, A.M., et al. (2020). CFTR Modulator Therapy Enhances Peripheral Blood Monocyte Contributions to Immune Responses in People With Cystic Fibrosis. Frontiers in pharmacology 11, 1219. 10.3389/fphar.2020.01219.

32. Quon, B.S., Dai, D.L.Y., Hollander, Z., Ng, R.T., Tebbutt, S.J., Man, S.F.P., Wilcox, P.G., and Sin, D.D. (2016). Discovery of novel plasma protein biomarkers to predict imminent cystic fibrosis pulmonary exacerbations using multiple reaction monitoring mass spectrometry. Thorax 71, 216–222. 10.1136/thoraxjnl-2014-206710.

33. Roberts, J.M., Dai, D.L.Y., Hollander, Z., Ng, R.T., Tebbutt, S.J., Wilcox, P.G., Sin, D.D., and Quon, B.S. (2018). Multiple reaction monitoring mass spectrometry to identify novel plasma protein biomarkers of treatment response in cystic fibrosis pulmonary exacerbations. Journal of cystic fibrosis : official journal of the European Cystic Fibrosis Society 17, 333–340. 10.1016/j.jcf.2017.10.013.

34. Graham, B.I.M., Harris, J.K., Zemanick, E.T., and Wagner, B.D. (2023). Integrating airway microbiome and blood proteomics data to identify multi-omic networks associated with response to pulmonary infection. The microbe 1. 10.1016/j.microb.2023.100023.

35. Dong, K., Moon, K.-M., Chen, V., Ng, R., Foster, L.J., Tebbutt, S.J., and Quon, B.S. (2019). Identification of novel blood biomarkers of treatment response in cystic fibrosis pulmonary exacerbations by label-free quantitative proteomics. Sci Rep 9, 17126. 10.1038/s41598-019-53759-1.

36. Benabdelkamel, H., Alamri, H., Okla, M., Masood, A., Abdel Jabar, M., Alanazi, I.O., Alfadda, A.A., Nizami, I., Dasouki, M., and Abdel Rahman, A.M. (2020). Serum-Based Proteomics Profiling in Adult Patients with Cystic Fibrosis. International Journal of Molecular Sciences 21, 7415. 10.3390/ijms21197415.

37. DeBoer, E.M., Kroehl, M.E., Wagner, B.D., Accurso, F.J., Harris, J.K., Lynch, D.A., Sagel, S.D., and Deterding, R.R. (2017). Proteomic profiling identifies novel circulating markers associated with bronchiectasis in cystic fibrosis. Proteomics. Clinical applications 11. 10.1002/prca.201600147.

38. Hendry, J., Elborn, J.S., Nixon, L., Shale, D.J., and Webb, A.K. (1999). Cystic fibrosis: inflammatory response to infection with Burkholderia cepacia and Pseudomonas aeruginosa. The European respiratory journal 14, 435–438. 10.1034/j.1399-3003.1999.14b32.x.

39. Sagel, S.D., Wagner, B.D., Ziady, A., Kelley, T., Clancy, J.P., Narvaez-Rivas, M., Pilewski, J., Joseloff, E., Sha, W., and Zelnick, L., et al. (2020). Utilizing centralized biorepository samples for biomarkers of cystic fibrosis lung disease severity. Journal of cystic fibrosis : official journal of the European Cystic Fibrosis Society 19, 632–640. 10.1016/j.jcf.2019.12.007.

40. Quon, B.S., Ngan, D.A., Wilcox, P.G., Man, S.F.P., and Sin, D.D. (2014). Plasma sCD14 as a biomarker to predict pulmonary exacerbations in cystic fibrosis. PLoS ONE 9, e89341. 10.1371/journal.pone.0089341.

41. Bene, Z., Fejes, Z., Macek, M., Amaral, M.D., Balogh, I., and Nagy, B. (2020). Laboratory biomarkers for lung disease severity and progression in cystic fibrosis. Clinica chimica acta; international journal of clinical chemistry 508, 277–286. 10.1016/j.cca.2020.05.015.

42. Charro, N., Hood, B.L., Faria, D., Pacheco, P., Azevedo, P., Lopes, C., Almeida, A.B. de, Couto, F.M., Conrads, T.P., and Penque, D. (2011). Serum proteomics signature of cystic fibrosis patients: a complementary 2-DE and LC-MS/MS approach. Journal of proteomics 74, 110–126. 10.1016/j.jprot.2010.10.001.

43. Kuroda, K., Ishii, K., Mihara, Y., Kawanoue, N., Wake, H., Mori, S., Yoshida, M., Nishibori, M., and Morimatsu, H. (2021). Histidine-rich glycoprotein as a prognostic biomarker for sepsis. Sci Rep 11, 10223. 10.1038/s41598-021-89555-z.

44. Elborn, J.S., Horsley, A., MacGregor, G., Bilton, D., Grosswald, R., Ahuja, S., and Springman, E.B. (2017). Phase I Studies of Acebilustat: Biomarker Response and Safety in Patients with Cystic Fibrosis. Clinical and translational science 10, 28–34. 10.1111/cts.12428.

45. Allen, J.R., Ge, L., Huang, Y., Brauer, R., Parimon, T., Cassel, S.L., Sutterwala, F.S., and Chen, P. (2018). TIMP-1 Promotes the Immune Response in Influenza-Induced Acute Lung Injury. Lung 196, 737–743. 10.1007/s00408-018-0154-2.

46. Jain, R., Baines, A., Khan, U., Wagner, B.D., and Sagel, S.D. (2021). Evaluation of airway and circulating inflammatory biomarkers for cystic fibrosis drug development. Journal of cystic fibrosis : official journal of the European Cystic Fibrosis Society 20, 50–56. 10.1016/j.jcf.2020.06.017.

47. Gray, R.D., Imrie, M., Boyd, A.C., Porteous, D., Innes, J.A., and Greening, A.P. (2010). Sputum and serum calprotectin are useful biomarkers during CF exacerbation. Journal of cystic fibrosis : official journal of the European Cystic Fibrosis Society 9, 193–198. 10.1016/j.jcf.2010.01.005.

48. Lee, S.J., Jeong, J.H., Heo, M., Ju, S., Yoo, J.-W., Jeong, Y.Y., and Lee, J.D. (2022). Serum Fibrinogen as a Biomarker for Disease Severity and Exacerbation in Patients with Non-Cystic Fibrosis Bronchiectasis. Journal of clinical medicine 11. 10.3390/jcm11143948.

49. Causer, A.J., Shute, J.K., Cummings, M.H., Shepherd, A.I., Gruet, M., Costello, J.T., Bailey, S., Lindley, M., Pearson, C., and Connett, G., et al. (2020). Circulating biomarkers of antioxidant status and oxidative stress in people with cystic fibrosis: A systematic review and meta-analysis. Redox Biology 32, 101436. 10.1016/j.redox.2020.101436.

50. Nixon, L.S., Yung, B., Bell, S.C., Elborn, J.S., and Shale, D.J. (1998). Circulating immunoreactive interleukin-6 in cystic fibrosis. American journal of respiratory and critical care medicine 157, 1764–1769. 10.1164/ajrccm.157.6.9704086.

51. Hanssens, L.S., Cellauro, S., Duchateau, J., and Casimir, G.J. (2021). Immunoglobulin G: A useful outcome marker in the follow-up of cystic fibrosis patients? Immunity, Inflammation and Disease 9, 608–614. 10.1002/iid3.426.

52. Bulloch, M.N., Hanna, C., and Giovane, R. (2017). Lumacaftor/ivacaftor, a novel agent for the treatment of cystic fibrosis patients who are homozygous for the F580del CFTR mutation. Expert review of clinical pharmacology 10, 1055–1072. 10.1080/17512433.2017.1378094.

53. Lepissier, A., Bonnel, A.S., Wizla, N., Weiss, L., Mittaine, M., Bessaci, K., Kerem, E., Houdouin, V., Reix, P., and Marguet, C., et al. (2023). Moving the Dial on Airway Inflammation in Response to Trikafta in Adolescents with Cystic Fibrosis. American journal of respiratory and critical care medicine 207, 792–795. 10.1164/rccm.202210-1938LE.

54. Ernst, G., Dantas, E., Sabatté, J., Caro, F., Salvado, A., Grynblat, P., and Geffner, J. (2015). Histidine-rich glycoprotein and idiopathic pulmonary fibrosis. Respiratory medicine 109, 1589– 1591. 10.1016/j.rmed.2015.10.010.

55. Wake, H., Mori, S., Liu, K., Morioka, Y., Teshigawara, K., Sakaguchi, M., Kuroda, K., Gao, Y., Takahashi, H., and Ohtsuka, A., et al. (2016). Histidine-Rich Glycoprotein Prevents Septic Lethality through Regulation of Immunothrombosis and Inflammation. EBioMedicine 9, 180–194. 10.1016/j.ebiom.2016.06.003.

56. Terao, K., Wake, H., Adachi, N., Liu, K., Teshigawara, K., Takahashi, H., Mori, S., and Nishibori, M. (2018). Histidine-Rich Glycoprotein Suppresses Hyperinflammatory Responses of Lung in a Severe Acute Pancreatitis Mouse Model. Pancreas 47, 1156–1164. 10.1097/MPA.0000000000001153.

57. Elborn, J.S., Konstan, M.W., Taylor-Cousar, J.L., Fajac, I., Horsley, A., Sutharsan, S., Aaron, S.D., Daines, C.L., Uluer, A., and Downey, D.G., et al. (2021). Empire-CF study: A phase 2 clinical trial of leukotriene A4 hydrolase inhibitor acebilustat in adult subjects with cystic fibrosis. Journal of cystic fibrosis : official journal of the European Cystic Fibrosis Society 20, 1026–1034. 10.1016/j.jcf.2021.08.007.

58. Snelgrove, R.J., Jackson, P.L., Hardison, M.T., Noerager, B.D., Kinloch, A., Gaggar, A., Shastry, S., Rowe, S.M., Shim, Y.M., and Hussell, T., et al. (2010). A critical role for LTA4H in limiting chronic pulmonary neutrophilic inflammation. Science (New York, N.Y.) 330, 90–94. 10.1126/science.1190594.

59. Li, L., and Somerset, S. (2014). Digestive system dysfunction in cystic fibrosis: challenges for nutrition therapy. Digestive and liver disease : official journal of the Italian Society of Gastroenterology and the Italian Association for the Study of the Liver 46, 865–874. 10.1016/j.dld.2014.06.011.

60. Peretti, N., Roy, C.C., Drouin, E., Seidman, E., Brochu, P., Casimir, G., and Levy, E. (2006). Abnormal intracellular lipid processing contributes to fat malabsorption in cystic fibrosis patients. American journal of physiology. Gastrointestinal and liver physiology 290, G609–15. 10.1152/ajpgi.00332.2005.

61. Thompson, S.J., Sargsyan, A., Lee, S.-A., Yuen, J.J., Cai, J., Smalling, R., Ghyselinck, N., Mark, M., Blaner, W.S., and Graham, T.E. (2017). Hepatocytes Are the Principal Source of Circulating RBP4 in Mice. Diabetes 66, 58–63. 10.2337/db16-0286.

62. Fedders, R., Muenzner, M., Weber, P., Sommerfeld, M., Knauer, M., Kedziora, S., Kast, N., Heidenreich, S., Raila, J., and Weger, S., et al. (2018). Liver-secreted RBP4 does not impair glucose homeostasis in mice. The Journal of biological chemistry 293, 15269–15276. 10.1074/jbc.RA118.004294.

63. Oh, H.S.-H., Rutledge, J., Nachun, D., Pálovics, R., Abiose, O., Moran-Losada, P., Channappa, D., Urey, D.Y., Kim, K., and Sung, Y.J., et al. (2023). Organ aging signatures in the plasma proteome track health and disease. Nature 624, 164–172. 10.1038/s41586-023-06802-1.

64. Steinhoff, J.S., Lass, A., and Schupp, M. (2021). Biological Functions of RBP4 and Its Relevance for Human Diseases. Frontiers in physiology 12, 659977. 10.3389/fphys.2021.659977.

65. Jin, Q., Chen, Y., Lou, Y., and He, X. (2013). Low Serum retinol-binding protein-4 levels in acute exacerbations of chronic obstructive pulmonary disease at intensive care unit admission is a predictor of mortality in elderly patients. Journal of inflammation (London, England) 10, 31. 10.1186/1476-9255-10-31.

66. Liu, X., Yang, M., Li, J., Liu, H., Dong, Y., Zheng, J., and Huang, Y. (2024). Identification of CFH and FHL2 as biomarkers for idiopathic pulmonary fibrosis. Frontiers in medicine 11, 1363643. 10.3389/fmed.2024.1363643.

67. Saraswat, M., Joenväärä, S., Tohmola, T., Sutinen, E., Vartiainen, V., Koli, K., Myllärniemi, M., and Renkonen, R. (2020). Label-free plasma proteomics identifies haptoglobin-related protein as candidate marker of idiopathic pulmonary fibrosis and dysregulation of complement and oxidative pathways. Sci Rep 10, 7787. 10.1038/s41598-020-64759-x.

68. Taguchi, A., Politi, K., Pitteri, S.J., Lockwood, W.W., Faça, V.M., Kelly-Spratt, K., Wong, C.-H., Zhang, Q., Chin, A., and Park, K.-S., et al. (2011). Lung cancer signatures in plasma based on proteome profiling of mouse tumor models. Cancer Cell 20, 289–299. 10.1016/j.ccr.2011.08.007.

69. Wikoff, W.R., Hanash, S., DeFelice, B., Miyamoto, S., Barnett, M., Zhao, Y., Goodman, G., Feng, Z., Gandara, D., and Fiehn, O., et al. (2015). Diacetylspermine Is a Novel Prediagnostic Serum Biomarker for Non-Small-Cell Lung Cancer and Has Additive Performance With Pro-Surfactant Protein B. Journal of clinical oncology : official journal of the American Society of Clinical Oncology 33, 3880–3886. 10.1200/JCO.2015.61.7779.

70. Sin, D.D., Tammemagi, C.M., Lam, S., Barnett, M.J., Duan, X., Tam, A., Auman, H., Feng, Z., Goodman, G.E., and Hanash, S., et al. (2013). Pro-surfactant protein B as a biomarker for lung cancer prediction. Journal of clinical oncology : official journal of the American Society of Clinical Oncology 31, 4536–4543. 10.1200/JCO.2013.50.6105.

71. Doyle, I.R., Bersten, A.D., and Nicholas, T.E. (1997). Surfactant proteins-A and -B are elevated in plasma of patients with acute respiratory failure. American journal of respiratory and critical care medicine 156, 1217–1229. 10.1164/ajrccm.156.4.9603061.

72. Robin, M., Dong, P., Hermans, C., Bernard, A., Bersten, A.D., and Doyle, I.R. (2002). Serum levels of CC16, SP-A and SP-B reflect tobacco-smoke exposure in asymptomatic subjects. The European respiratory journal 20, 1152–1161. 10.1183/09031936.02.02042001.

73. Probst, C.K., Montesi, S.B., Medoff, B.D., Shea, B.S., and Knipe, R.S. (2020). Vascular permeability in the fibrotic lung. European Respiratory Journal 56. 10.1183/13993003.00100-2019.

74. Um, S.J., Lam, S., Coxson, H., Man, S.F.P., and Sin, D.D. (2013). Budesonide/formoterol enhances the expression of pro Surfactant Protein-B in lungs of COPD patients. PLoS ONE 8, e83881. 10.1371/journal.pone.0083881.

75. Perez-Riverol, Y., Bai, J., Bandla, C., García-Seisdedos, D., Hewapathirana, S., Kamatchinathan, S., Kundu, D.J., Prakash, A., Frericks-Zipper, A., and Eisenacher, M., et al. (2022). The PRIDE database resources in 2022: a hub for mass spectrometry-based proteomics evidences. Nucleic acids research 50, D543–D552. 10.1093/nar/gkab1038.

76. Graeber, S.Y., Vitzthum, C., Pallenberg, S.T., Naehrlich, L., Stahl, M., Rohrbach, A., Drescher, M., Minso, R., Ringshausen, F.C., and Rueckes-Nilges, C., et al. (2022). Effects of Elexacaftor/Tezacaftor/Ivacaftor Therapy on CFTR Function in Patients with Cystic Fibrosis and One or Two F508del Alleles. American journal of respiratory and critical care medicine 205, 540–549. 10.1164/rccm.202110-2249OC.

77. Rappsilber, J., Ishihama, Y., and Mann, M. (2003). Stop and go extraction tips for matrix-assisted laser desorption/ionization, nanoelectrospray, and LC/MS sample pretreatment in proteomics. Analytical chemistry 75, 663–670. 10.1021/ac026117i.

78. Pekayvaz, K., Leunig, A., Kaiser, R., Joppich, M., Brambs, S., Janjic, A., Popp, O., Nixdorf, D., Fumagalli, V., and Schmidt, N., et al. (2022). Protective immune trajectories in early viral containment of non-pneumonic SARS-CoV-2 infection. Nature communications 13, 1018. 10.1038/s41467-022-28508-0.

79. Keshishian, H., Burgess, M.W., Gillette, M.A., Mertins, P., Clauser, K.R., Mani, D.R., Kuhn, E.W., Farrell, L.A., Gerszten, R.E., and Carr, S.A. (2015). Multiplexed, Quantitative Workflow for Sensitive Biomarker Discovery in Plasma Yields Novel Candidates for Early Myocardial Injury. Molecular & Cellular Proteomics : MCP 14, 2375–2393. 10.1074/mcp.M114.046813.

80. Nordmeyer, S., Kraus, M., Ziehm, M., Kirchner, M., Schafstedde, M., Kelm, M., Niquet, S., Stephen, M.M., Baczko, I., and Knosalla, C., et al. (2023). Disease- and sex-specific differences in patients with heart valve disease: a proteome study. Life science alliance 6. 10.26508/lsa.202201411.

81. Tyanova, S., Temu, T., and Cox, J. (2016). The MaxQuant computational platform for mass spectrometry-based shotgun proteomics. Nat Protoc 11, 2301–2319. 10.1038/nprot.2016.136.

82. Gordon Smyth, Yifang Hu, Matthew Ritchie, Jeremy Silver, James Wettenhall, Davis McCarthy, Di Wu, Wei Shi, Belinda Phipson, AaronLun, Natalie Thorne, Alicia Oshlack, Carolynde Graaf, Yunshun Chen, Mette Langaas, EgilFerkingstad, Marcus Davy, Francois Pepin, Dongseok Choi (2017). limma (Bioconductor).

83. Subramanian, A., Tamayo, P., Mootha, V.K., Mukherjee, S., Ebert, B.L., Gillette, M.A., Paulovich, A., Pomeroy, S.L., Golub, T.R., and Lander, E.S., et al. (2005). Gene set enrichment analysis: a knowledge-based approach for interpreting genome-wide expression profiles. Proceedings of the National Academy of Sciences of the United States of America 102, 15545–15550. 10.1073/pnas.0506580102.

